# Synaptic Signatures and Disease Vulnerabilities of Layer 5 Pyramidal Neurons

**DOI:** 10.1101/2024.01.22.576602

**Authors:** Gabriele Marcassa, Dan Dascenco, Blanca Lorente-Echeverría, Danie Daaboul, Jeroen Vandensteen, Elke Leysen, Lucas Baltussen, Andrew J. M. Howden, Joris de Wit

## Abstract

Cortical layer 5 (L5) intratelencephalic (IT) and pyramidal tract (PT) neurons are embedded in distinct information processing pathways. The morphology, connectivity, electrophysiological properties, and role in behavior of these neurons have been extensively analyzed. However, the molecular composition of their synapses remains largely uncharacterized. Here, we dissect the protein composition of the excitatory postsynaptic compartment of L5 neurons in intact somatosensory circuits, using an optimized proximity biotinylation workflow with subsynaptic resolution. We find distinct synaptic signatures of L5 IT and PT neurons that are defined by proteins regulating synaptic organization and transmission, including cell-surface proteins (CSPs), neurotransmitter receptors and ion channels. In addition, we find a differential vulnerability to disease, with a marked enrichment of autism risk genes in the synaptic signature of L5 IT neurons compared to PT neurons. Our results align with human studies and suggest that the excitatory postsynaptic compartment of L5 IT neurons is notably susceptible in autism. Together, our analysis sheds light on the proteins that regulate synaptic organization and function of L5 neuron types and contribute to their susceptibility in disease. Our approach is versatile and can be broadly applied to other neuron types to create a protein-based, synaptic atlas of cortical circuits.

## Introduction

How different types of neurons become embedded in neural circuits, the basis of the brain’s capacity for information processing, is a major question in neuroscience. Cortical L5 pyramidal neurons (PNs) play a crucial role in information processing^1–3^. These neurons integrate feedforward sensory information converging on their basal dendritic compartment and feedback contextual information converging on their apical dendrites^4,5^. Their axons project to cortical and subcortical regions^6,7^. L5 PNs are thus at the heart of cortical input/output processing.

Two main populations of L5 PNs can broadly be distinguished: intratelencephalic (IT) and pyramidal tract (PT) L5 PNs, which differ in their morphology, electrophysiological properties and projection targets^6,8^. IT neurons are thin-tufted, regular spiking and intracortically projecting PNs mostly located in superficial L5 (L5a), whereas PT neurons are thick-tufted, bursting and subcortically projecting PNs mostly found in deeper L5 (L5b)^9,10^. Both L5 neuron types project to the striatum, however, they carry different types of information^7,11^. The two L5 PN types differ in input connectivity, which is best characterized for excitatory input from the thalamus. L5 IT neurons primarily receive input from higher-order thalamic nuclei, whereas PT neurons preferentially receive input from the primary thalamus^12–14^. Functionally, L5 IT and PT neurons are part of different cortical information processing pathways. Perception of tactile stimuli depends on activation of apical dendrites of L5 PT, but not IT, neurons in the somatosensory cortex^5^. In the motor cortex, L5 IT neurons are involved in preparation of movement, whereas PT neurons are important for initiation and execution^10,15^. In the visual cortex, PT neurons display higher direction selectivity index and prefer higher temporal frequencies than IT neurons, indicating that they integrate different visual information^16^. Differences in connectivity and function of the two L5 PN types may also relate to their differential vulnerability in neurodevelopmental and movement disorders^6^.

Despite the wealth of information on morphological and functional properties, the molecular composition of L5 PN synapses has not been characterized. Elucidating cell type-specific molecular synaptic signatures could contribute to our understanding of how L5 PN types are connected into circuits and how perturbations in this process might contribute to diseases, particularly those characterized by a strong synaptic component, such as neurodevelopmental disorders^17^. Proximity biotinylation uses a promiscuous biotin ligase to covalently attach a biotin molecule to nearby proteins within a short radius^18^ and offers an attractive approach to profile synaptic proteomes. This approach has been used to characterize the molecular composition of postsynaptic compartments from whole cortex and hippocampus^19^, and the subcellular proteome of astrocytes in the striatum^20^, as well as to profile the neuronal proteomes of major neuronal classes expressing a cytosolic biotin ligase^21–23^. However, profiling synaptic protein composition of genetically defined populations of cortical neurons is challenging, due to the small number of starting cells and correspondingly low amounts of input material.

To pave the way for such a systematic description of the synaptic protein composition of cortical neurons, we optimize a cell type-specific proximity biotinylation-based workflow using the engineered biotin ligase TurboID^24^. We fuse TurboID to two different excitatory postsynaptic scaffold proteins, PSD95 and Homer1, and demonstrate the high spatial resolution of this approach in L5 neurons *in vivo*. We then analyze the protein composition of the excitatory postsynaptic compartment of L5 IT and PT neurons in the somatosensory cortex. We find that the synaptic signatures of L5 IT and PT neurons are characterized by differential expression of CSPs, neurotransmitter receptors and ion channels. To assess disease vulnerability of the L5 IT and PT neuron synaptic signatures, we analyze the identified proteomes for disease association. We find a striking enrichment of ASD risk genes in the excitatory postsynaptic proteome of L5 IT neurons compared to PT neurons, suggesting that the excitatory postsynaptic compartment of L5 IT neurons is notably susceptible in ASD. Cross-referencing our results with human transcriptomics data found several matches in human L5 IT neurons. Our results fit in a framework of L5 IT neuron vulnerability in ASD^6^ and align with RNA sequencing studies of individuals with autism that highlight vulnerability of IT neurons^25–28^. Taken together, our analysis sheds light on the proteins that regulate synaptic organization and function of L5 PN types and contribute to their vulnerability in disease.

## Results

### Efficient, compartment-specific protein biotinylation in genetically identified neurons

To compare the molecular synaptic signatures of L5 IT and PT neurons in the somatosensory cortex, we first optimized a workflow to identify postsynaptic proteomes in a cell-type specific manner, with optimal capture of biotinylated proteins from a small population of neurons in a specific brain region. This optimized workflow is based on the high-efficiency biotin ligase TurboID (Fig. 1a), which has a reported ∼10-fold increase in *in vitro* activity compared to previously published biotin ligases^24^. To restrict the activity of TurboID to the excitatory postsynaptic density (PSD), we fused TurboID to the scaffold proteins PSD95 (PSD95.turboID) and Homer1^29^ (Homer1.turboID) (Fig. 1b). As a control, we generated a TurboID construct without any localization sequence (cytosolic.turboID) (Fig. 1b). To target TurboID to L5 PNs in a cell-type and region-specific manner, we packaged Cre-dependent TurboID expression constructs into adeno-associated viral vectors (AAV) and injected them in the primary somatosensory (S1) cortex of postnatal day (P)28 mouse lines expressing Cre recombinase in L5 PNs. A week after stereotactic injection, we subcutaneously injected the TurboID substrate biotin and collected brains for immunofluorescence or biochemical analysis. Tissue sections stained for HA-tagged TurboID and for biotinylated proteins using fluorescent streptavidin showed colocalizing signal restricted to S1 and concentrated in L5 (Fig. 1c). A control sample in which only biotin but no virus was injected (no.turboID) showed that background biotinylation was below detection levels compared to other conditions (Fig. 1c). High-resolution confocal imaging showed a diffuse pattern of biotinylated proteins throughout the neuronal cytoplasm in cytosolic.turboID samples (Fig. 1d). In PSD95.turboID and Homer1.turboID samples, streptavidin-labeled biotinylated proteins displayed a punctate pattern that colocalized with the excitatory postsynaptic markers Homer1 and PSD95, respectively, confirming TurboID activity in the excitatory postsynaptic compartment (Fig. 1d).

**Figure 1.**
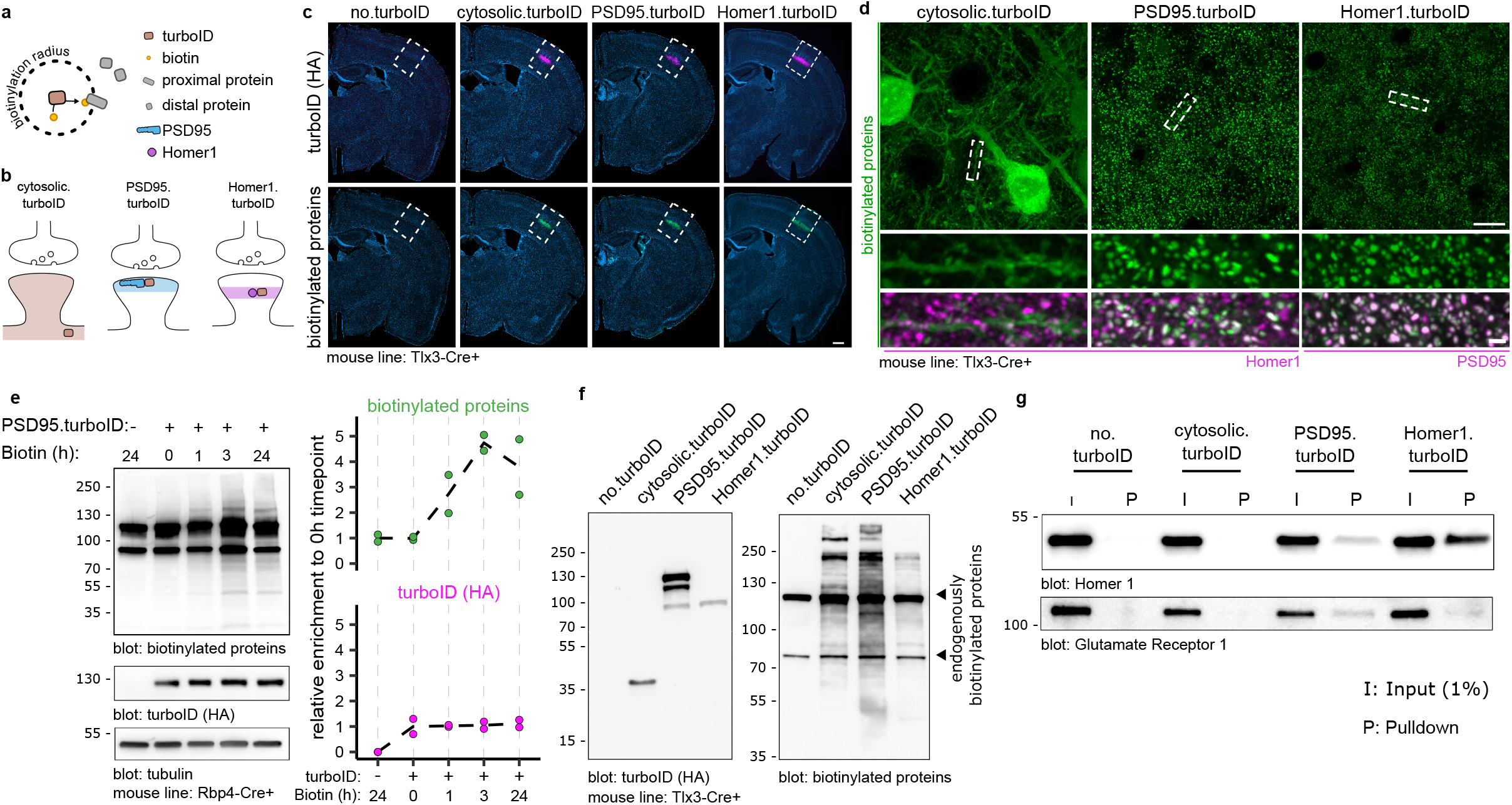
Fast and specific postsynaptic proximity biotinylation in genetically identified cortical layer 5 neurons. **a,** TurboID covalently binds a biotin molecule to nearby proteins. **b**, A cytosolic.turboID construct labels cytosolic proteins. Fusion of TurboID to PSD95 (PSD95.turboID) or Homer1 (Homer1.turboID) directs its enzymatic activity to distinct postsynaptic compartments. **c,** Whole-section imaging of Tlx3-Cre mice injected with Cre-dependent TurboID constructs labelled for HA-tagged TurboID enzyme and biotinylated proteins. Scalebar is 200 μm. **d,** High resolution imaging shows biotinylated protein patterns from the TurboID constructs. Homer1 and PSD95 immunostaining labels postsynapses. Scalebar is 10 μm and 1 μm in the cropped region. **e,** Western blot quantification shows high levels of biotinylated proteins in dissected somatosensory cortices after 3 hours of biotin injection. Tubulin is used as loading control (N = 2 mice per time point). **f,** Western blot for TurboID confirms expression of constructs at the expected size. Both TurboID constructs produce strong biotinylated protein signal across molecular weights compared to no.turboID sample, which shows only bands for endogenously biotinylated proteins. **g,** After streptavidin pulldown, postsynaptic marker proteins Gria1 and Homer1 are only detected in PSD95.turboID and in Homer1.turboID samples.

To control for potential artefacts resulting from fusion protein expression, we analyzed spine density and morphology in TurboID-expressing samples using the Supernova labelling system, which allows for sparse yet bright GFP expression in genetically identified neurons^30^ (Extended Data Fig. 1). In initial experiments, we noticed that higher PSD95.turboID viral titers resulted in an increase in mushroom-like spines (Extended Data Fig. 1a). We therefore adjusted viral titers and performed dendritic spine reconstruction and classification, and found no difference in spine density or morphology across conditions at lower titers (Extended Data Fig. 1b-d).

To optimize *in vivo* biotinylation, we expressed PSD95.turboID in L5 PNs in somatosensory cortex and provided biotin for different durations before collecting tissue for biochemical analysis. Western blot quantification showed a maximum, 4-fold increase in biotinylation levels compared to control conditions at 3h following biotin injection (Fig. 1e). We therefore selected 3h post-biotin injection for subsequent experiments to achieve maximum biotinylation while limiting off-target labelling^31^.

We then compared *in vivo* expression and activity of the different TurboID constructs. HA immunoblot showed that the TurboID constructs run at the expected size of 35kDa for cytosolic.turboID, ∼135 kDa for PSD95.turboID, and ∼100 kDa for Homer1.turboID (Fig. 1f). Immunoblotting for biotinylated proteins showed two bands in the no.turboID condition, likely corresponding to endogenously biotinylated proteins^32^, while cytosolic, PSD95.turboID and Homer1.turboID produced biotinylation across all molecular weights (Fig. 1f). Streptavidin pulldown from the same lysate detected signal for postsynaptic proteins Gria1 and Homer1 in the PSD95.turboID and Homer1.turboID samples, but not in the other conditions (Fig. 1g). Together, these results show that our proximity biotinylation strategy reliably targets the excitatory postsynaptic compartment of genetically identified L5 PNs in a specific cortical region and efficiently yields biotinylated proteins in a short timeframe.

### Identification of excitatory postsynaptic proteins in cortical L5 neurons with subsynaptic resolution

The PSD95 and Homer1 scaffold proteins are core components of the PSD but reside in different locations^29^ (Fig. 2a). To determine the extent of coverage of PSD proteins and analyze potential differences in proteins identified by the two fusion proteins, we performed a comparison of PSD95.turboID and Homer1.turboID in L5 neurons. To this end, we injected L5 PN-specific Tlx3_PL56-Cre mice with TurboID-expressing AAVs at P28 (Fig. 2b). One week later, mice were injected with biotin for 3h followed by tissue dissection and collection of the somatosensory cortex. Negative control samples (no.turboID) were prepared with mice injected only with biotin (Fig. 2a), to exclude endogenously biotinylated proteins and contaminants from the analysis. Four mice were used per replicate. Biotinylated proteins were purified from tissue lysates using streptavidin-coupled beads, eluted, and digested to peptides by trypsin and identified by label-free liquid chromatography-tandem mass spectrometry (LC-MS/MS) run in data-independent acquisition (DIA) mode (Fig. 2b) (see Methods). Protein identification and quantification performed with DIA-NN software^33^ yielded more than 4000 different proteins among all samples, which showed consistent intensity distribution at the whole sample level (Extended Data Fig. 2a).

**Figure 2.**
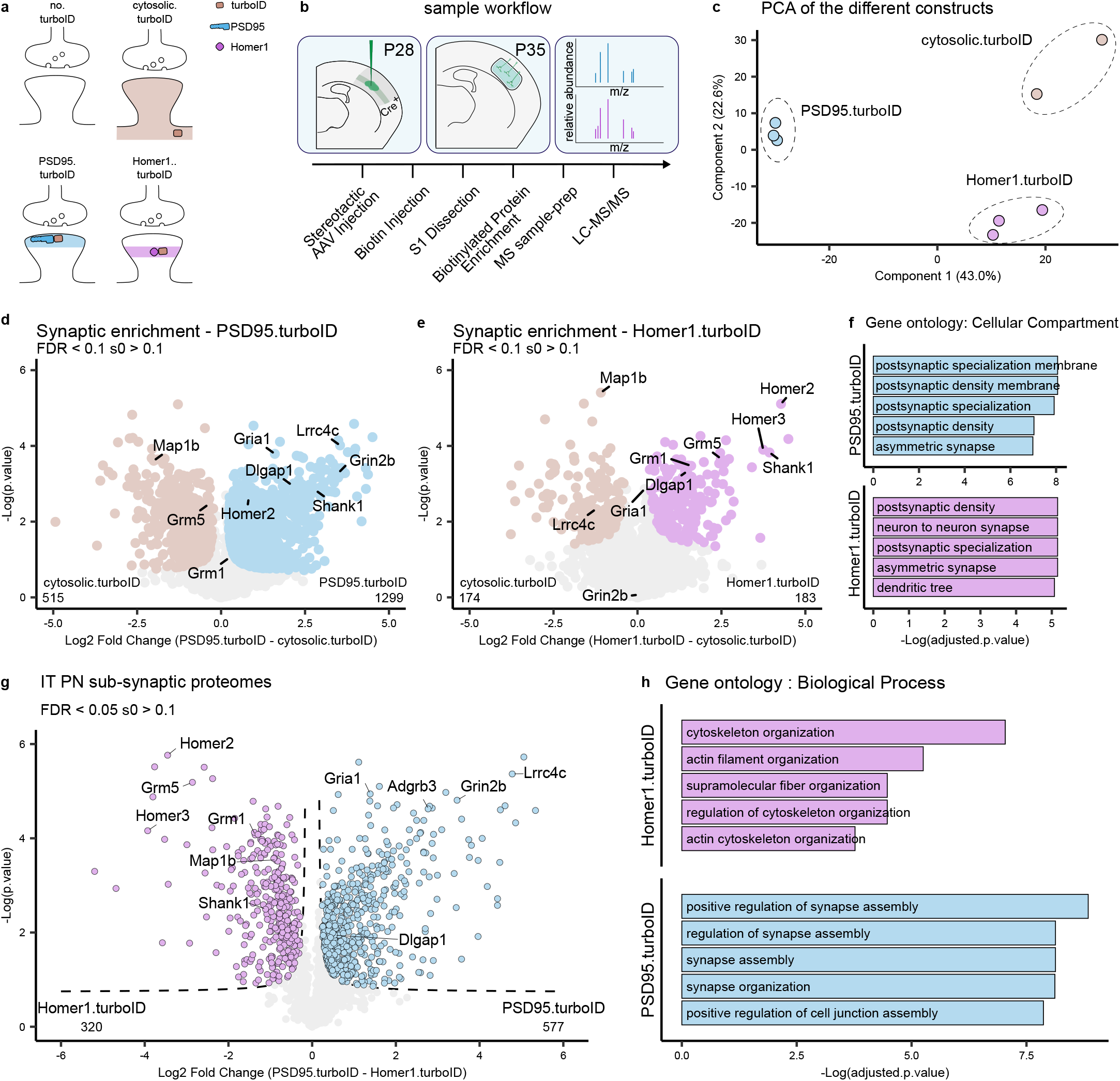
Identification of L5 cortical neuron proteins with subsynaptic resolution. **a,** Representation of the different conditions tested and the expected pattern of biotinylated proteins at the synapse. **b,** Mice are injected with AAVs in the somatosensory cortex at P28. At P35, biotin is provided 3 hours before collection of somatodendritic material from somatosensory cortices. Biotinylated proteins are enriched by streptavidin pulldown and prepared for MS analysis. **c,** PCA analysis shows replicates clustering by TurboID construct. **d,** Relative protein enrichment between PSD95.turboID and cytosolic.turboID. Significant proteins (unpaired t-test with permutation-based FDR correction at 10%, s0 > 0.1) are labelled, core synaptic proteins are highlighted. **e,** Relative protein enrichment between Homer1.turboID and cytosolic.turboID. Significant proteins (unpaired t-test with permutation-based FDR correction at 10%, s0 > 0.1) are labelled, core synaptic proteins are highlighted. **f,** Gene ontology analysis of synaptic proteins in PSD95.turboID and Homer1.turboID samples. **d, e, g**, Numbers of differentially expressed proteins for each probe are indicated in bottom left and right corners of Volcano plots. **g,** Relative protein enrichment by PSD95.turboID and Homer1.turboID. Significant proteins (unpaired t-test with permutation-based FDR correction at 5%, s0 > 0.1) are labelled based on TurboID construct. **h,** Gene ontology of differentially expressed proteins in PSD95.turboID vs. Homer1.turboID samples.

As expected, we detected high levels of endogenously biotinylated proteins in all samples (Extended Data Fig. 2b). Postsynaptic proteins on the other hand showed selective enrichment in the PSD95.turboID and Homer1.turboID replicates and depletion in the no.turboID samples (Extended Data Fig. 2c). We used an approach similar to a previously published ratiometric analysis^24^ to create a list of proteins detected in the no.turboID condition and selected contaminants to be excluded from the analysis (Extended Data Fig. 2d) (see Methods). PCA analysis showed replicates clustering together by the TurboID construct expressed (Fig. 2c).

To determine the extent of PSD protein coverage of the two constructs, we compared PSD95.turboID and Homer1.turboID with the cytosolic.turboID samples and selected proteins with a false discovery rate (FDR) < 0.1 and s0 > 0.1 (Fig. 2d, e). Both PSD95.turboID and Homer1.turboID fusion proteins identified PSD constituents such as Shank1 and Dlgap1, although the number of synaptic proteins identified with Homer1.turboID was lower compared to PSD95.turboID (Fig. 2d, e), but (Fig. 2d, e). Gene Ontology (GO) analysis (cellular compartment) confirmed PSD localization of synaptically enriched proteins for PSD95.turboID and Homer1.turboID (Fig. 2f). To further characterize the synaptic proteins identified by PSD95.turboID and Homer1.turboID, we queried the synaptic protein database published in Sorokina, Mclean, Croning et al.^34^ (from here on referred to as the ‘synapse proteome’ database). This database lists the synaptic proteins detected in proteomic studies performed using various approaches in the last 20 years and collects and integrates information with several external databases. More than 80% of the proteins identified in the PSD95.turboID and Homer1.turboID samples were present in this database (Extended Data Fig 2e and Supplementary Table 1). Cross-referencing with the curated synaptic protein database SYNGO^35^ showed an overlap of 21% of identified proteins in PSD95.turboID and 29% in Homer1.turboID, due to its relatively limited coverage (Extended Data Fig 2e).

To analyze potential differences in the synaptic proteins detected in the PSD95.turboID and Homer1.turboID samples, we performed a differential expression analysis (Fig. 2g). PSD95 interactors, including neurotransmitter receptors such as the NMDA receptor subunit Grin2b^36^ and CSPs such as Lrrc4c^37^, were enriched in the PSD95.turboID sample (Fig. 2g). Homer1 interactors, such as the postsynaptic metabotropic glutamate receptors Grm1 and Grm5^38^, were enriched in the Homer1.turboID sample (Fig. 2g). GO analysis (biological process) showed more synaptic terms for PSD95.turboID compared to Homer1.turboID (Fig. 2h). Together, these results show that our optimized proximity biotinylation strategy combined with DIA-MS identifies postsynaptic proteins with subsynaptic resolution in genetically identified cortical neurons. Based on its broader synaptic protein coverage and stronger enrichment of neurotransmitter receptors and CSPs (Fig. 2d-h) that are important for synaptic function and organization, we chose to use PSD95.turboID for subsequent experiments.

### Identification of excitatory postsynaptic proteins in cortical L5 IT and PT neurons

We next applied our approach to characterize the postsynaptic protein composition of L5 IT and PT neurons. To obtain genetic access to these neurons, we used the Tlx3_PL56-Cre and Sim1_KJ18-Cre mouse lines^5,16,39,40^. The projection patterns, as well as morphological and physiological properties of Tlx3-Cre+ and Sim1-Cre+ neurons in the cortex have been found to represent thin-tufted L5a IT and thick-tufted L5b PT neurons, respectively^5,7^. Imaging of tissue sections from Tlx3-Cre and Sim1-Cre mice injected with an AAV expressing Cre-dependent GFP confirmed the segregation of labeled Tlx3-Cre+ and Sim1-Cre+ cell bodies to superficial and deep L5, respectively (Fig. 3a, b). We injected P28 Tlx3-Cre or Sim1-Cre mice with cytosolic.turboID or PSD95.turboID AAVs and collected somatosensory cortex at P35 following 3 hours of biotin injection (Fig. 3c, d). We reduced the number of animals per replicate to two, obtaining around 1mg of total dissected tissue. Before biotinylated protein purification, TurboID expression and successful biotinylation was confirmed by western blot (Extended Data Fig. 3a). After LC-MS/MS, protein identification and quantification yielded more than 4500 different proteins among all samples, which showed consistent intensity distribution at the whole sample level (Extended Data Fig. 3b). Consistent with our previous experiments, endogenously biotinylated proteins were detected in all samples, while postsynaptic proteins showed highest enrichment in the PSD95.turboID samples (Extended Data Fig. 3c, d). We again created a list of contaminants to be excluded from the analysis based on the no.turboID samples (Extended Data Fig. 3e, f) (see Methods). PCA analysis showed replicates clustering together by condition with little separation relative to the Cre line used (Fig. 3e).

**Figure 3.**
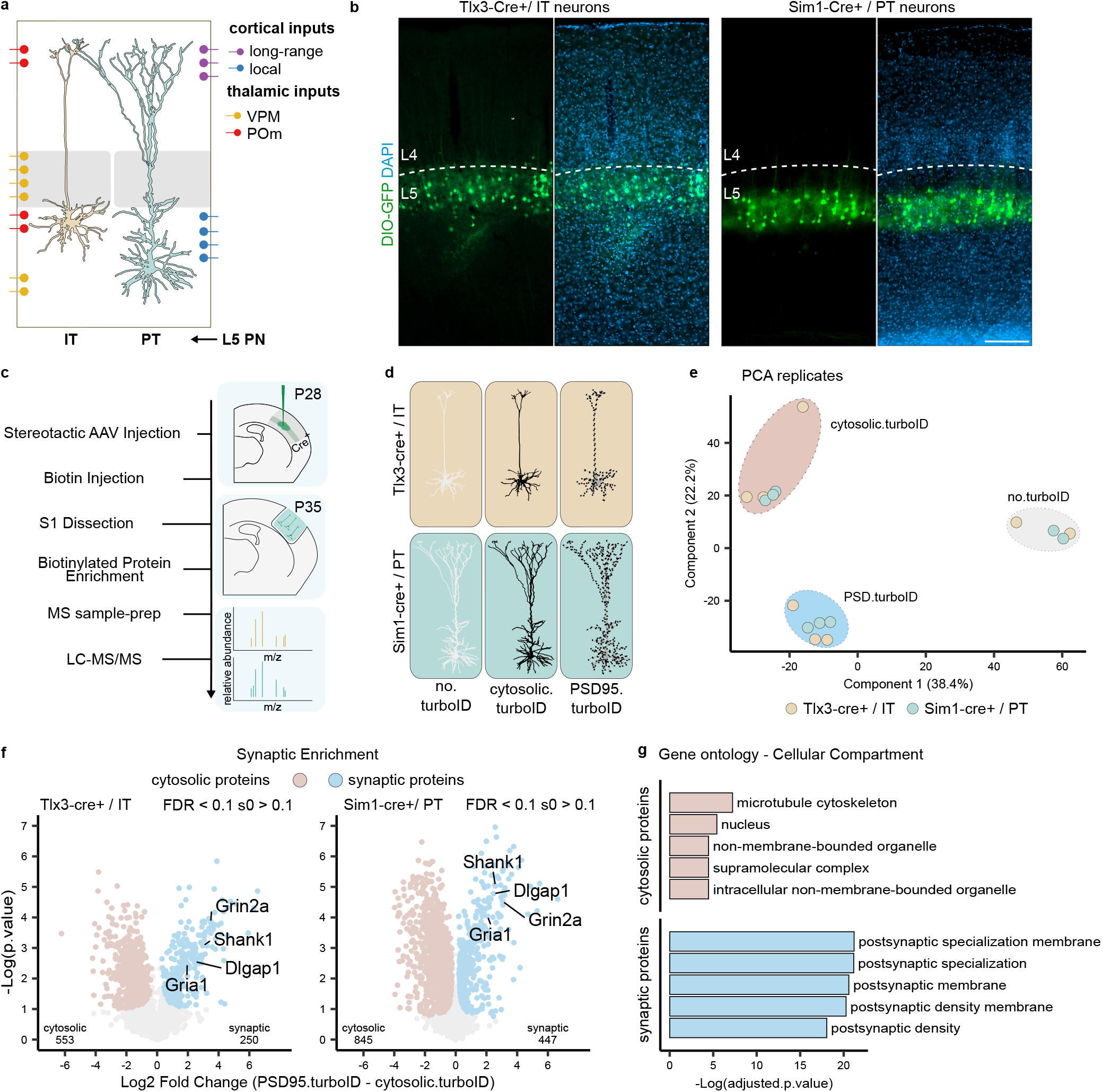
Identification of the postsynaptic proteome of layer 5 neurons. **a,** Representation of L5 IT and PT neuron position and morphology in the somatosensory cortex with overview of input localization from other local and long-range excitatory neurons. (Adapted from ^8^). **b,** AAV-mediated Cre-dependent GFP expression in Tlx3-Cre+ and Sim1-Cre+ mouse lines recapitulates soma position for L5 IT and PT neurons, respectively. Layer 4/5 borders were identified by density of DAPI-positive nuclei. Scalebar is 200 μm. **c,** Workflow: mice are injected with AAVs in the somatosensory cortex at P28. At P35, biotin is provided 3 hours before collection of somatosensory cortices. Biotinylated proteins are enriched by streptavidin pulldown and prepared for MS analysis. **d,** MS replicates were collected from Tlx3-Cre+ and Sim1-Cre+ mice injected with no.turboID, cytosolic.turboID or PSD95.turboID. **e,** PCA analysis shows replicates clustering by condition but not by Cre line. **f,** Statistical analysis of cytosolic.turboID and PSD95.turboID samples for each Cre line allows for synaptic protein enrichment. Core constituents of the excitatory postsynapse are annotated. Numbers of identified cytosolic and synaptic proteins for Tlx3-Cre+ and Sim1-Cre+ lines are indicated in bottom left and right corners of each graph. Unpaired t-test with permutation-based FDR correction at 10%, s0 > 0.1. **g,** Gene ontology analysis of the cytosolic and synaptic proteins from (f) showing cellular localization of identified proteins.

Comparison of the cytosolic.turboID and PSD95.turboID conditions for each Cre line showed core constituents of the excitatory postsynapse, such as glutamate receptors and scaffolding proteins, to be synaptically enriched in both L5 Tlx3-Cre+/IT and Sim1-Cre+/PT samples (Fig. 3f). Postsynaptic localization was confirmed by GO analysis (cellular compartment) (Fig. 3h), showing that we can reliably identify synaptic proteins even when halving the number of animals used.

### Synaptic signatures of cortical L5 IT and PT neurons

To map the synaptic signatures of L5 Tlx3-Cre+/IT and Sim1-Cre+/PT neurons and analyze potential differences between IT and PT neurons, we next performed a differential expression analysis of the synaptically enriched proteins of the two cell types. As expected, most synaptic proteins did not differ significantly between L5 IT and PT neurons, but 135 proteins were differentially expressed between Tlx3-Cre+/IT (93 proteins) and Sim1-Cre+/PT (42 proteins) neurons (Fig. 4a and Supplementary Table 2). We first analyzed the function of these proteins using several GO databases^41^. Differentially expressed proteins showed strong postsynaptic localization (Extended Data Fig. 4a). ‘Synapse organization’ and ‘regulation of postsynaptic neurotransmitter receptor activity’ were among the most represented GO terms in the Biological Process category (Fig. 4b). The Molecular Function category showed enrichment mostly in terms related to PSD interactions (Extended Data Fig. 4b). Of the identified synaptic proteins in L5 Tlx3-Cre+/IT and Sim1-Cre+/PT neurons, 84% were represented in the synapse proteome database (Extended Data Fig. 4c) and 44% in SYNGO (Extended Data Fig. 4d, e).

**Figure 4.**
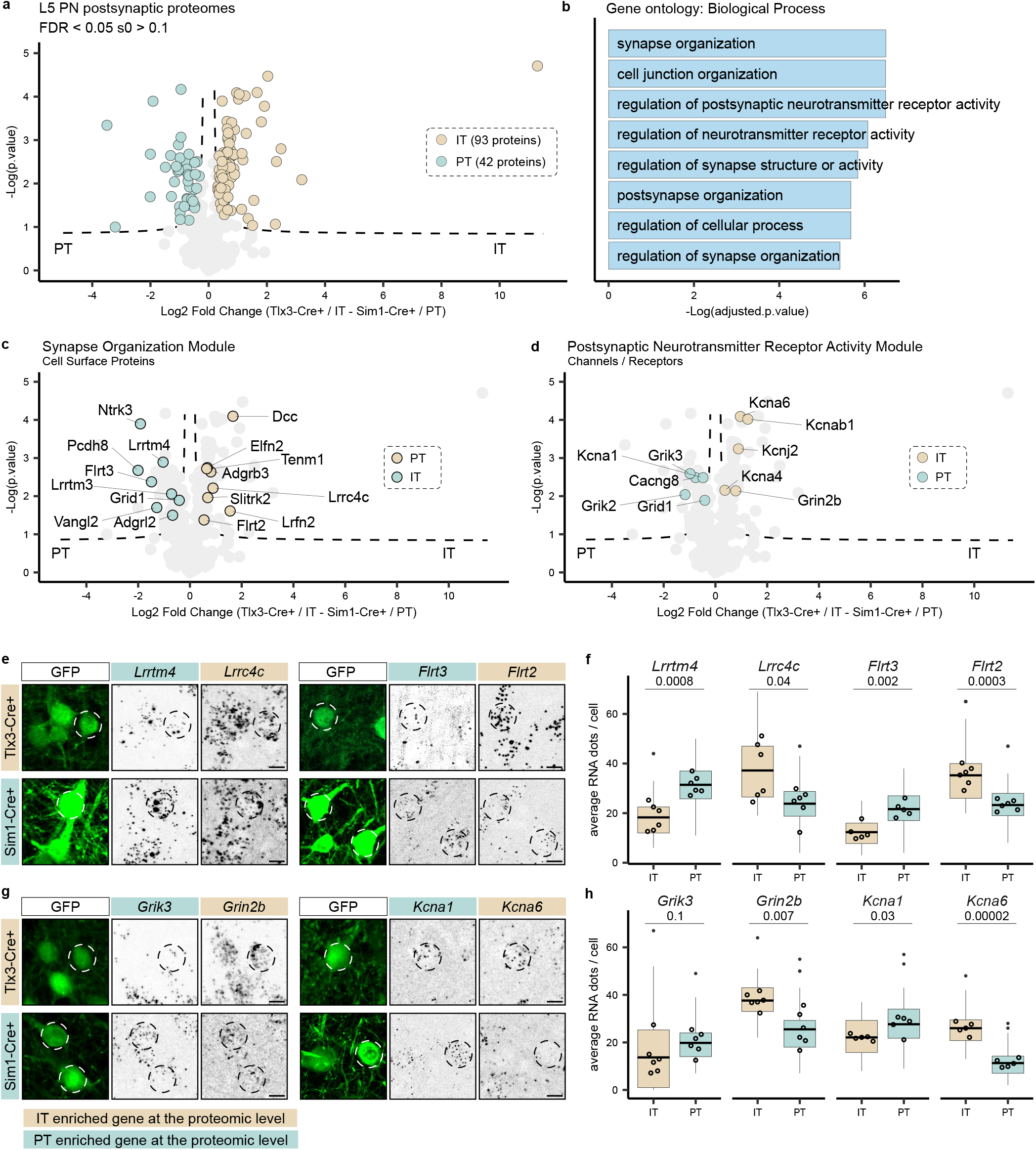
Molecular synaptic signatures of two populations of layer 5 neurons. **a,** Relative protein enrichment in L5 IT and PT neurons. Significant proteins (unpaired t-test with permutation-based FDR correction at 5%, s0 > 0.1) are labelled according to Cre line (93 in Tlx3-Cre/IT and 42 in Sim1-Cre/PT neurons). **b,** Biological Process gene ontology analysis of differentially expressed proteins. **c,** CSPs are annotated to represent the ’synapse organization’ module. **d,** Ion channels and neurotransmitter receptors are annotated to represent the ‘postsynaptic neurotransmitter receptor activity’ module. **e, g,** smFISH analysis of candidate genes in L5 IT and PT neurons labelled by Cre-dependent GFP expression in Tlx3-Cre and Sim1-Cre mice, respectively. Cells that have been quantified are indicated by dotted circles. Scale bars 10 μm. **f, h,** Quantification of RNA expression (average number of RNA puncta per cell). N = 5 or 6 mice per Cre line, 8 cells per mouse are quantified. Boxplots (line across the box describes the mean) show distribution of single cells, empty dots display the average of each biological replicate. Statistical analysis is performed as an unpaired t-test, alpha = 0.05, p-value is reported under transcript name.

We next explored the proteins in the ‘synapse organization’ and ‘regulation of postsynaptic neurotransmitter receptor activity’ modules, as these are likely to contribute to connectivity and physiological properties of Tlx3-Cre+/L5 IT and Sim1-Cre+/PT neurons. To do so, we started from the proteins annotated in each term of the Biological Process category (Fig. 4b and Extended Data Fig. 4f) and integrated these results with PANTHER^42^ and Uniprot^43^ information to assign a function to the differentially expressed proteins (see Supplementary Table 2). In the ‘synapse organization module’, CSPs were the most abundant protein class (Fig. 4c). Strikingly, extracellular leucine-rich repeat (LRR) proteins (Elfn2, Flrt2, Flrt3, Lrrtm3, Lrrtm4, Lrrc4c/Ngl-1, Ntrk3/Trkc, Lrfn2/Salm1, Slitrk2) were the most represented protein family among the differentially expressed CSPs in L5 IT and PT neurons (Fig. 4c). Interestingly, Flrt2 and Flrt3 belong to the same LRR subfamily but showed differential enrichment in IT and PT neuron synaptic signatures (Fig. 4c). LRR proteins control precise wiring and confer synaptic properties across species^44–46^, but their role in cortical circuit connectivity and function remains largely unexplored.

In the ‘postsynaptic neurotransmitter receptor activity’ module, neurotransmitter receptors and channels were the most enriched protein classes (Fig. 4d). Among differentially expressed receptors, kainate receptors (Grik2, Grik3) and their auxiliary subunit Neto1 showed enrichment in the L5 PT neuron synaptic signature (Fig. 4d and Extended Data Fig. 4f). Kainate receptors have been observed to be enriched in deep cortical layers of the mouse brain using immunohistochemistry^47^ and receptor autoradiography^48,49^, but an enrichment in L5 PT neurons has not been reported to our knowledge. The NMDA receptor subunit Grin2b was enriched in IT neurons, whereas potassium and calcium ion channels showed different subunits specifically enriched in one of the two L5 neuron types (Fig. 4d and Extended Data Fig. 4f).

Due to limitations in availability of validated neuroscience antibodies^50^ and challenges in localizing synaptic proteins to specific neuron types in cortex, we decided to assess our proteomics results using an orthogonal approach measuring cell-type specific gene expression. While it has been reported that mRNA and protein rarely match at absolute levels, it has also been shown that mRNA can predict proteomic differences at a global level^51–53^. We selected four differentially expressed proteins from the ‘synapse organization’ and from the ‘regulation of postsynaptic neurotransmitter receptor activity’ modules (Fig. 4e-h). We injected AAVs expressing Cre-dependent GFP in Tlx3-Cre or Sim1-Cre mice and performed single-molecule Fluorescence In Situ Hybridization (smFISH) experiments (Fig. 4e, g). Quantification of mRNA puncta in GFP+ cells showed significant agreement between cell type enrichment of the selected synaptic proteins and their corresponding transcripts (Fig. 4f, h), except for *Grik3*, which showed a bimodal distribution in Tlx3-Cre+/IT neurons with cells expressing high levels of mRNA and cells expressing none (Extended Data Fig. 4g). These results show that for these selected proteins at least, cell type-specific differences in synaptic protein enrichment correlate with differences at the transcript level. In conclusion, these results show that the synaptic signatures of L5 IT and PT neurons are defined by differential expression of CSPs, neurotransmitter receptors and ion channels.

### Differential vulnerability of L5 IT and PT synaptic signatures in neurodevelopmental disorders

Neurodevelopmental disorders are associated with altered synaptic function and development^17,54–56^. We next tested whether the synaptic signatures of Tlx3-Cre+/IT and Sim1-Cre+/PT neurons might provide insight in disease vulnerability of these two L5 neuron types. We first ran the differentially expressed synaptic proteins in Tlx3-Cre+/IT and Sim1-Cre+/PT neurons (Fig. 4a) through the GeneTrek database^57^, which provides association of genes with neurodevelopmental disorders (Fig. 5a). Based on several disease association databases, genes are given a ’high confidence’ or ’candidate’ score. The synaptic signatures of both L5 IT and PT neurons showed a high proportion of genes in each of these two categories (Fig. 5a). Next, we queried the synapse proteome database for disease association and graphed the results for L5 IT and PT neurons, normalized to the total number of synaptic proteins identified in each neuron type (Fig. 5b and Supplementary Table 2). Overall, neurodevelopmental disorders showed a higher percentage of enriched genes compared to neurodegenerative diseases such as Parkinson’s or Alzheimer’s in IT and PT neurons (Fig. 5b and Supplementary Table 2). The synaptic signature of PT neurons showed a larger fraction of genes associated with schizophrenia and psychotic disorder, whereas that of IT neurons showed a larger fraction of genes associated with ASD and intellectual disability (Fig. 5b and Supplementary Table 2).

**Figure 5.**
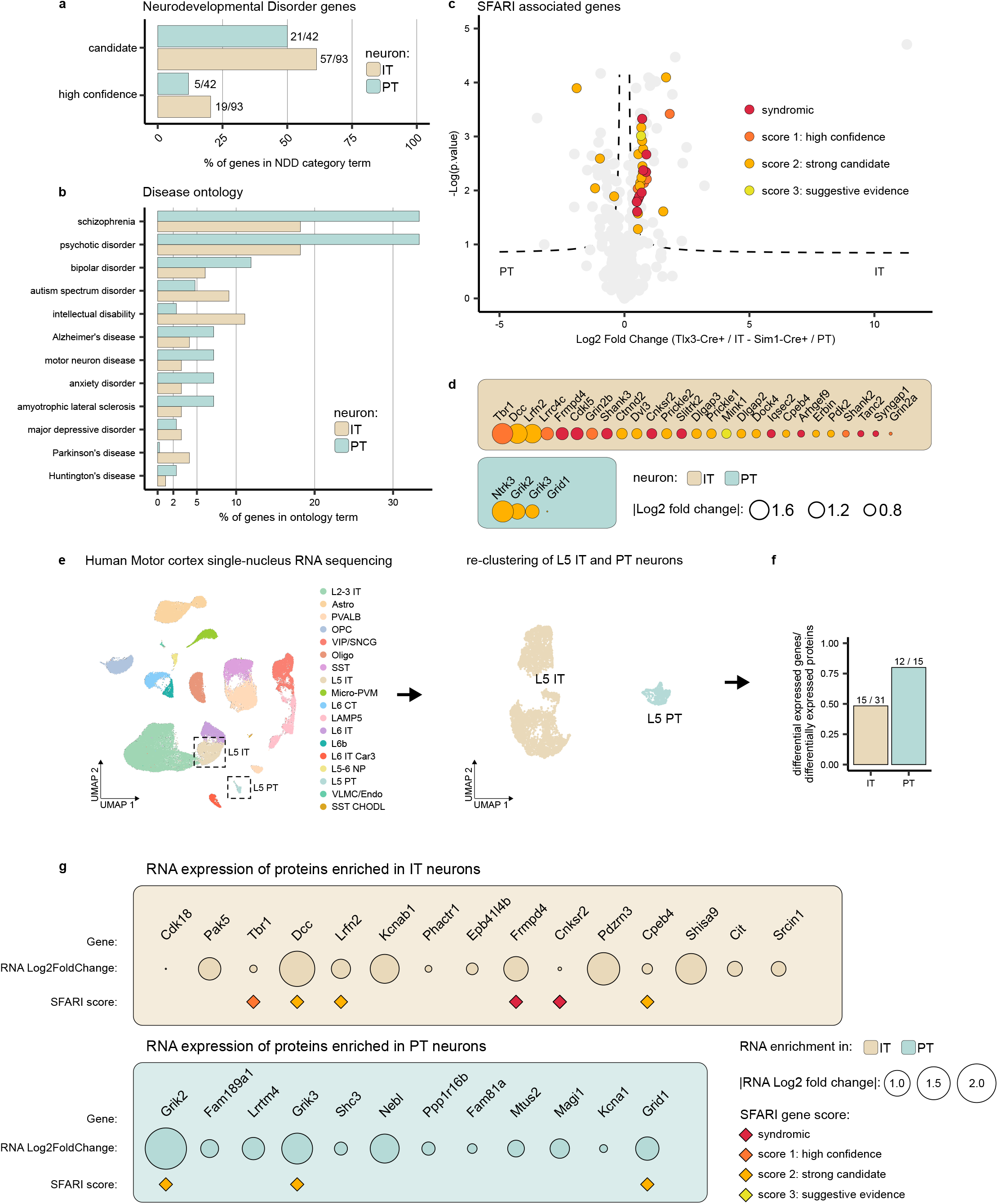
Differential disease vulnerability of layer 5 neuron molecular synaptic signatures. **a,** Percentage of synaptic genes per cell type present in the ’high confidence’ or ’candidate’ neurodevelopmental disorder lists based on the Genetrek database published in Leblond et al., 2021^32^. **b,** Percentage of synaptic genes per cell type present in each disorder term based on the synapse proteome database^6^. **c,** Synaptic proteins in L5 IT and PT neurons with an ASD risk score annotation in the SFARI gene database^33^ are indicated on the volcano plot showing protein distribution. **d,** ASD proteins identified in the SFARI database^33^ listed per cell type and ranked by protein log fold change. **e,** L5 IT and PT neurons were extracted from a published human motor cortex snRNA-Seq data^61^ and re-clustered. **f,** Differentially expressed proteins in mouse where compared to differentially expressed genes in human motor cortex. Of the 93 differentially expressed proteins detected in IT mouse neurons and 42 proteins in PT mouse neurons, 31 and 15 genes respectively showed differential expression also in human samples. **g,** Mouse proteins that show matching gene differential expression in human are shown. SFARI scores are annotated.

The SFARI gene database^58^ has a curated, up to date annotation of ASD genes scored by clinical and scientific evidence. Based on the larger fraction of genes associated with ASD and intellectual disability in IT neurons (Fig. 5b), we therefore cross-referenced the synaptic signatures of Tlx3-Cre+/IT and Sim1-Cre+/PT neurons with the SFARI gene database. This analysis revealed a marked enrichment of ASD risk genes in the synaptic signature of L5 IT neurons compared to that of PT neurons (Fig. 5c). ASD genes with the highest risk score annotation (‘syndromic’ and ‘high confidence’) in the SFARI database, such as Frmpd4, Cdkl5, Shank3, Cnksr2 and Slitrk2, were all found in the L5 IT synaptic signature (Fig. 5c, d), suggesting that the excitatory postsynaptic compartment of L5 IT neurons is notably susceptible in ASD.

Finally, to translate the significance of our findings to human neurons, we extracted and re-clustered L5 IT and PT neuron transcriptomes from a publicly available human motor cortex single-nucleus RNA sequencing data set^59^ and calculated differential gene expression between IT and PT neurons (Fig. 5e) (See Methods). We next tested whether differentially expressed synaptic proteins in mouse L5 IT or PT neurons were also differentially expressed at the RNA level in human L5 IT and PT neurons. Of 93 differentially expressed synaptic proteins in mouse IT neurons, we found 31 to be differentially expressed at the RNA level in human neurons; 48% of which showed matching enrichment in human L5 IT neurons (Fig. 5f). Of 42 differentially expressed synaptic proteins in mouse PT neurons, 15 were differentially expressed at the RNA level in human neurons; 80% of which showed matching enrichment in human L5 PT neurons (Fig. 5f). Moreover, 6 out of 15 IT genes enriched both at the mouse protein and human RNA level were annotated in the SFARI database (Fig. 5g). Together, these results show that L5 IT and PT synaptic signatures display differential vulnerability to neurodevelopmental disorders such as ASD, and that these results can be translated to human neurons.

## Discussion

Here, we characterize the synaptic signatures of two closely related L5 cortical neuron types from a specific brain region, using an optimized proximity biotinylation workflow with subsynaptic resolution. CSPs, neurotransmitter receptors and ion channels with a role in synapse organization and synaptic transmission define the synaptic signatures of L5 IT and PT neurons, which are differentially vulnerable in disease. The synaptic signature of L5 PT neurons shows a larger fraction of genes associated with schizophrenia and psychotic disorder, whereas the IT synaptic signature displays a marked enrichment of ASD risk genes.

We find that our approach is sufficiently sensitive to identify proteins with subsynaptic resolution from L5 cortical neurons *in vivo*. PSD95.turboID and Homer1.turboID both identified PSD proteins, but interactors of each fusion protein showed differential enrichment in the corresponding condition. CSPs and neurotransmitter receptors were enriched in the PSD95.turboID samples, whereas the postsynaptic metabotropic glutamate receptors Grm1 and Grm5 were enriched in the Homer1.turboID samples, demonstrating the high spatial resolution of this approach in a small population of genetically identified neurons. We focused on PSD95.turboID for our experiments, because of the relevance of the identified proteins for synapse function and organization, but depending on the application, other parts of the synapse can be targeted using this approach.

The cell type-specific synaptic signatures we identify represent promising targets for future investigation into regulators of synaptic properties, physiology and connectivity of L5 IT and PT neurons. Especially striking in this context are the LRR family proteins, which dominate the CSPs in the L5 Tlx3-Cre+/IT and Sim1-Cre+/PT neuron synapse organization modules. LRR proteins are important regulators of connectivity and synaptic properties^60–62^. Input-specific combinations of postsynaptic LRR proteins define synaptic structure and function in hippocampal PNs^63^. Analogous to hippocampal PNs^63–65^, LRR proteins may differentially distribute across the dendritic tree of cortical PNs to regulate connectivity, structure and function of specific inputs. While our approach does not provide spatial information about synaptic protein distribution in L5 PN dendrites, this remains an important point that will need to be explored in further studies.

L5 Tlx3-Cre+/IT neurons displayed a higher number of differentially expressed synaptic proteins than Sim1-Cre+/PT neurons. PCA and sample level intensity analysis exclude differences in labelling efficiency between the two cell types. The higher number of differentially expressed synaptic proteins in L5 IT neurons compared to PT neurons could reflect differences in synapse number between the two neuron types or differences in type of excitatory input. Compared to L5 PT neurons, L5 IT neurons have a less extensive dendritic arborization, which would argue against an increased synapse number in IT neurons. The two L5 PN neuron types have excitatory input in common but also show cell type-specific differences in input connectivity patterns^12^. First-order input from the ventral posteromedial (VPM) thalamic nucleus preferentially, but not exclusively, targets L5 PT neurons, whereas higher-order thalamic input from the posteromedial (POm) nucleus is concentrated on L5 IT neurons^12–14,66,67^. It will be of interest to determine whether the synaptic signatures of L5 IT and PT neurons contribute to differences in thalamic input connectivity.

We find that the synaptic signatures of L5 Tlx3-Cre+/IT and Sim1-Cre+/PT neurons display differential vulnerability to distinct neurodevelopmental disorders. L5 PT neurons showed a larger fraction of synaptic proteins associated with schizophrenia, whereas IT neurons showed a larger fraction of synaptic proteins associated with ASD. Cross-referencing with the SFARI gene database revealed a striking enrichment of syndromic and high confidence ASD risk genes in the synaptic signature of L5 IT neurons. These findings are in line with RNA sequencing studies that find IT neurons to be the most affected cell types in ASD cortex^25,26,68^ and suggest altered intracortical processing in ASD^27,69^. Our synaptic compartment-specific proteome data indicates that compared to Sim1-Cre+/PT neurons, excitatory input of Tlx3-Cre+/IT neurons is particularly vulnerable in ASD. Moreover, comparing the differentially expressed proteins with single-nuclei RNA sequencing data from human motor cortex samples shows good agreement between mouse proteins and human genes in IT and PT neurons. Considering the differences in excitatory input connectivity between L5 IT and PT neurons, it is tempting to speculate that higher-order thalamic input on IT neurons is affected in ASD. Brain imaging studies have indeed observed altered thalamocortical functional connectivity in individuals with autism^70–73^. Whether this involves projections from first- or higher-order thalamic nuclei to specific cortical neuron types is unknown.

In addition to formulating hypotheses on L5 IT and PT connectivity and function, the synaptic signatures we identify here can also be used to explore specific hypotheses on protein function. An interesting candidate from our analysis is Tbr1, which is enriched in the Tlx3-Cre+/IT synaptic signature and a high confidence ASD risk gene. Tbr1 is a transcription factor but shows a dendritic distribution in pyramidal neurons and localizes to synaptic fractions in biochemical analysis^74^. Tbr1 binds the synaptic scaffold protein CASK, which translocates to the nucleus^75^. Possibly, PSD-associated Tbr1 shuttles between synapse and nucleus, where it can regulate expression levels of Grin2b^76^, which we also find to be enriched in the L5 IT synaptic signature. Another interesting candidate we identified is Pelo, which is highly enriched in the synaptic signature of Tlx3-Cre+/IT neurons. Pelo rescues stalled ribosomes and triggers degradation of damaged mRNA in a process termed No-Go Decay^77,78^ but its role at synapses is unknown.

To obtain these results, we optimized every step of the proximity biotinylation workflow, resulting in the identification of thousands of proteins starting from single, genetically identified L5 neuron populations in the somatosensory cortex. We found that a single subcutaneous biotin injection 3 hours before sample collection is sufficient to produce strong protein biotinylation in the brain. Previous *in vivo* studies have relied on multiple injections of biotin over several days^19,22,23,79^. As prolonged labeling with biotin increases the number of proteins identified at the expense of specificity^24^, minimizing the biotinylation window is important for the analysis of compartment-specific protein composition. Short biotinylation windows also open new possibilities, for example to study how the composition of synaptic proteomes changes upon neuronal activity. For similar reasons, we limited the duration of expression of TurboID constructs to 1 week (compared to multiple weeks or months in previous studies^19,23,79–81^), as low levels of protein biotinylation using endogenous biotin can occur while TurboID is expressed.

A possible limitation of our approach is the use of Cre driver lines to target IT and PT neurons. Although we selected the Tlx3-Cre and Sim1-Cre lines based on their thorough characterization for L5 IT and PT neurons in somatosensory cortex^5^, this approach is inherently limited to the selectivity of the Cre line used. An interesting alternative approach to explore would be to rely on connectivity patterns to target IT and PT neurons, using retrograde AAV. Another possible limitation of our approach is the use of PSD95 to target TurboID to the postsynaptic compartment. We carefully titrated our viral vectors to control PSD95 and TurboID expression levels and analyzed spine density and morphology *in vivo* to verify the absence of overexpression artefacts. This strategy is suitable for mature neurons but carries a risk of driving precocious postsynaptic maturation in developing neurons. Since we are targeting a small and defined population of cortical neurons in a specific brain region, efficient transduction is a requirement for obtaining sufficient amounts of protein for analysis. CRISPR/Cas9-mediated knock-in^80,82^ might be an alternative. This approach avoids the overexpression of exogenous proteins, however, the low percentage of successful recombination makes it challenging to target single neuronal populations in specific cortical regions^80,82^. A possible way to avoid modifying expression of synaptic proteins, while retaining short duration of TurboID expression and high efficiency of labeling, might be the use of recently published PSD95 binders^83^, which are reported not to alter PSD95 interactions.

In conclusion, our analysis of cell type- and compartment-specific protein composition sets a foundation for the systematical description of cortical circuit connectivity at the molecular synaptic level. As our approach enables the comparison of the synaptic proteome of single neuronal populations, and mouse Cre driver lines for cortical neuronal cell types are increasingly available^40,84^, systematic application of this workflow could create a protein-based, synaptic atlas of the cortex. Integration of synaptic signature information with RNA sequencing and electrophysiology data will be useful for computational modeling efforts and contribute to our understanding of circuit connectivity and function.

## Materials and Methods

### Marcassa et al. Synaptic Signatures and Disease Vulnerabilities of Layer 5 Pyramidal Neurons Animals

All animal experiments were conducted according to the KU Leuven ethical guidelines and approved by the KU Leuven Ethical Committee for Animal Experimentation. Mice were maintained in a specific pathogen-free facility under standard housing conditions with continuous access to food and water. Mice used in the study were 1–5-weeks old and were maintained on 14 h light, 10 h dark light cycle from 7 to 21 h. Wild-type (WT) C57BL/6J mice were obtained from JAX. Rpb4.cre and Tlx3.cre lines were obtained from GENSAT.

Sim1.cre was obtained from Dr. Charles Gerfen (National Institute of Mental Health, USA). Genotypes were regularly checked by PCR analysis. Heterozygous cre-positive males were breed to Bl6 females. For euthanasia, animals were either anesthetized with isoflurane, and decapitated or injected with an irreversible dose of ketamine–xylazine.

Cell Lines. HEK293T-17 human embryonic kidney cells were obtained from American Type Culture Collection (ATCC) cat# CRL-11268. HEK293T-17 cells were grown in Dulbecco’s modified Eagle’s medium (DMEM; Invitrogen) supplemented with 10% fetal bovine serum (FBS; Invitrogen) and penicillin/streptomycin (Invitrogen).

Plasmids TurboID sequence was a gift from Alice Ting (Addgene plasmid #107169). Mouse PSD95 sequence was a gift from Gary Bassell (Addgene plasmid #102949). TurboID was fused at the C-terminus of PSD95 o Homer1 and inserted into a cre-dependent AAV expression vector under the synapsin promoter by Gibson assembly (NEB). Plasmids will be deposited at plasmids.eu.

AAV purification High titer AAV production and purification was carried out as previously reported 1. Briefly, 6 plates of 70% confluent HEK cells were transfected with 20ug of pDelta F6 plasmid, 10ug of RepCap 2/9 plasmid and 10ug AAV genome plasmid per plate using PEI transfection. 3 days later cells were collected by scraping and centrifugation. AAV particles in the supernatant were precipitated by PEG and added to cell lysates. Lysates were treated by Benzonase nuclease to remove cellular DNA. Lysates were loaded onto a iodixanol gradient (60%, 40%, 25%, 15%) and ultracentrifuged 1:40 hours at 50.000 RPM at 12C. The 40% fraction containing the purified AAV was carefully collected and desalted and concentrated on a 4ml Amicon column (Sigma). AAV purity was tested by silver staining using the ProteoSilver™ Silver Stain Kit (Sigma PROTSIL1-1KT). AAV titer was determined by qPCR to normalize titers between different viral vector batches using LightCycler 480 SYBR Green I Master (Roche 04707516001). Primer for qPCR targeting the synapsin promoter were as follows: FWD tgataggggatgcgcaatttgg REV gtgcaagtgggttttaggacca.

Stereotactic injections P28 mice were anesthetized with 5% isoflurane and Duratears was applied to the eyes to prevent them from drying out. Mice were placed in a mouse stereotact (KOPF). During the rest of the procedure 2.5% isoflurane was constantly administered. After shaving and disinfecting the mouse’s head, local anesthesia was administered by a subcutaneous injection with 100 µl lidocaine (xylocain 1%). An incision was made on the skin to reveal the skull. A hole was drilled at coordinates for somatosensory cortex and 200nl of AAV mix injected using a Nanoject III (Drummond) though a beveled capillary at 2 nl/s. After a 5 min recovery, the capillary was pulled out at ∼0.1 mm/5 s. The incision was stitched with surgical glue (Millpledge Veterinary). After 6 h, their health was examined and mice were injected with 0.1 mg/kg buprenorphine. Two injections per hemisphere were performed for biochemistry experiments to cover a larger area of the somatosensory cortex and recover more material (x: 3.2, y: 0.3 and 1.5 from bregma, z: 0.8 from the surface of the brain). One injection per hemisphere was performed for every other experiment (x: 3.2, y: 0.9 from bregma, z: 0.8 from the surface of the brain). Cytoplasmic.turboID was injected at a ∼1.6*10^12 GC/mL, PSD95.turboID was injected at a ∼3*10^11 GC/mL, Homer1.turboID was injected at a ∼5*10^11 GC/mL, cre-dependent GFP was injected at ∼10^12 GC/mL.

Biotinylated protein collection One week after AAV injection, mice were injected subcutaneously with 1ml of 2mg/mL biotin in 1X PBS, adjusting the pH to 7.4 to increase biotin solubility. 3 hours later mice were anaesthetized by isoflurane and brains collected in ice cold HBSS. 500um coronal brain sections were produced using an ice-cold brain matrix (AgnThos 69-2165-1), and somatosensory cortices dissected under a fluorescence stereomicroscope using cre-dependent GFP signal expressed by infected cells and morphological landmarks. For experiment in Figure 2, tissue from L4 and L5 around GFP was dissected. For experiments in Figure 3 and 4, the whole cortical column around GFP was dissected. Tissue pieces from each mouse were collected in a separate tube, flash-frozen in liquid nitrogen and stored at - 80C until needed. For protein extraction and pulldown, we extensively optimized published protocols. Frozen tissue was slowly thawed on ice and tissue from 4 animals (Figure2) or 2 animals (Figure 3 and 4) was combined in a single tube as a replicate. Tissue was homogenized in 500ul 1% SDS RIPA buffer (50mM Tris pH 8, 150mM NaCl, 1% SDS, 0.5% Sodium Deoxycholate, 1% Triton-X 100, 1X protease inhibitors) using a small glass Dounce homogenizer. Lysates were sonicated 3x30s with 30s rest on ice in between. Lysates were boiled at 95C for 5 minutes to dissociate PSD and transsynaptic complexes and diluted to 0.5% SDS with ice cold SDS-free RIPA buffer as reported in 2. After 1 hour incubation at 4C with rotation, lysates were cleared by ultracentrifugation at 100.000G for 30 minutes at 4C and supernatant collected. BCA assay was used to quantify protein concentration. 10ug protein input was used for Western blot experiments in Figure 1F and Figure 1G. 200 ug protein input was used for pulldown for western blots in Figure 1H. Around 1000ug protein input was used for pulldown for MS. 5ul of streptavidin magnetic beads (Pierce) per 100ug of protein input was used for every condition. Beads were washed 3 times in 0.5% SDS RIPA buffer using a magnetic rack and incubated overnight at 4C with rotation with protein input in 1ml of 0.5% SDS RIPA buffer final volume. The day after beads were washed 3x with 0.5% SDS RIPA buffer, 1X with 1M KCl, 1x with 0.1M Na2CO3, 1x with 2M Urea in 50mM Tris-HCl ph 8. MS sample preparation steps are described below, for western blot analysis beads were further washed 3x with 0.5% SDS RIPA buffer and proteins eluted in 1X protein loading buffer supplemented with 10mM DTT and 2mM biotin 5 mins at 95C.

Western blot After boiling 5 minutes at 95C, samples were loaded on 4-20% or 7.5% polyacrylamide gels and run at 180V. Proteins were transferred to 0.2um nitrocellulose membrane using semi-dry transfer (Biorad) using the mixed-molecular weight program. Membranes were blocked in 5% milk in TBS-T buffer (25mM Tris-base pH 7.5, 300mM NaCl, 0.05% Tween-20) 1h at RT. Primary and secondary antibodies were diluted in 5% milk in TBS-T. Primary antibodies were incubated O/N at 4C, secondary antibodies were incubated 1h at RT. When blotting for biotinylated proteins, HRP-conjugated streptavidin was diluted 1:500 in 5% BSA to avoid binding to the biotin present in the milk. We obtain best results in streptavidin signal while blocking membranes in milk, however we noticed that some powdered milk gave higher background than other. Blocking in 5% BSA is an alternative. After primary and secondary antibodies membranes were washed 5 times with TBS-T and once with 1X PBS before development at ImageQuant 800 (Cytiva) using SuperSignal West Pico PLUS Chemiluminescent Substrate (Thermo Fisher 34577).

MS sample preparation Beads were washed 2 more times with 2M Urea in 50mM Tris-HCl pH 8. Proteins were eluted from the beads and digested to peptides using S-Trap kit (Protify) following manufacturer instructions. Beads were resuspended in 92 ul of 5% SDS 50 mM TEAB pH 8, boiled 5 minutes at 95C and let cool down. Eluates were reduced with 4 ul of a 120 mM TCEP buffer at 55 degrees for 15 mins. Eluates were alkylated with 4 ul of 500 mM MMTS in isopropanol at RT for 10 mins in the dark. Eluates were acidified with 10 ul of 27.5% phosphoric acid in dH2O. Using a magnetic rack to separate the beads, supernatants were collected in a different tube and diluted in 660ul of binding/wash buffer. Supernatant were loaded onto an S-trap column, centrifuged at 4000G 1 minute. Columns were washed 10 times with binding/wash buffer. Proteins were digested to peptides applying 1ug of trypsin diluted in 20ul of digestion buffer directly to the column and incubated overnight at 37C in a humidified oven. Peptides were sequentially eluted from the column using 40ul of elution buffer 1, 2 and 3 and dried using a speedVac before being shipped for LC-MS/MS.

Mass Spectrometry Peptides were analysed by data independent acquisition (DIA) mass spectrometry as described previously3,4. In summary, peptides were injected onto a nanoscale C18 reverse-phase chromatography system (UltiMate 3000 RSLC nano, Thermo Scientific) and electrosprayed into an Orbitrap Exploris 480 Mass Spectrometer (Thermo Fisher). For liquid chromatography the following buffers were used: buffer A (0.1% formic acid in Milli-Q water (v/v)) and buffer B (80% acetonitrile and 0.1% formic acid in Milli-Q water (v/v). Samples were loaded at 10 μL/min onto a trap column (100 μm × 2 cm, PepMap nanoViper C18 column, 5 μm, 100 Å, Thermo Scientific) equilibrated in 0.1% trifluoroacetic acid (TFA). The trap column was washed for 3 min at the same flow rate with 0.1% TFA then switched in-line with a Thermo Scientific, resolving C18 column (75 μm × 50 cm, PepMap RSLC C18 column, 2 μm, 100 Å). Peptides were eluted from the column at a constant flow rate of 300 nl/min with a linear gradient from 3% buffer B to 6% buffer B in 5 min, then from 6% buffer B to 35% buffer B in 115 min, and finally to 80% buffer B within 7 min. The column was then washed with 80% buffer B for 4 min and re-equilibrated in 3% buffer B for 15 min. Two blanks were run between each sample to reduce carry-over. The column was kept at a constant temperature of 50oC.

The data was acquired using an easy spray source operated in positive mode with spray voltage at 2.445 kV, and the ion transfer tube temperature at 250oC. The MS was operated in DIA mode. A scan cycle comprised a full MS scan (m/z range from 350-1650), with RF lens at 40%, AGC target set to custom, normalized AGC target at 300%, maximum injection time mode set to custom, maximum injection time at 20 ms, microscan set to 1 and source fragmentation disabled. MS survey scan was followed by MS/MS DIA scan events using the following parameters: multiplex ions set to false, collision energy mode set to stepped, collision energy type set to normalized, HCD collision energies set to 25.5, 27 and 30%, orbitrap resolution 30000, first mass 200, RF lens 40%, AGC target set to custom, normalized AGC target 3000%, microscan set to 1 and maximum injection time 55 ms. Data for both MS scan and MS/MS DIA scan events were acquired in profile mode.

MS data analysis RAW MS data were analysed by DIA-N-N version 1.8.15. Uniprot Mus musculus UP000000589 reference proteome was used for spectral library preparation. Precursor m/z range was set at 350 to 1650. Neural network classifier was set at double-pass mode. Every other parameter was kept at default. The resulting file was loaded onto Perseus for analysis6. A list of contaminants was created from the no.turboID samples. First, only proteins identified in all no.turboID samples were kept and ranked by intensity. Then, known PSD proteins were used to set a threshold defining true contaminants (Extended Data Fig. 2e). All keratins were considered contaminants and included in this list. Proteins in this list were removed from all other samples. Samples were log2 transformed and normalized by ‘substract median’ method. Only proteins present at least 3 times in at least one condition were kept (e.g. in all 3 Tlx3-Cre PSD95.turboID replicates) and missing values imputed with standard parameters. To enrich for synaptic proteins, we applied a two samples t-test with permutation-based FDR at 10% and s0 > 0.1 between the cytoplasmic.turboID and PSD95.turboID samples for each cre line. We then merged PSD95.turboID significant proteins from Tlx3-Cre and Sim1- Cre to create the ’synaptic protein list’ used for gene ontology and further comparison. To identify differentially expressed proteins between L5a and L5b neurons a t test was performed on the synaptic proteins with using an FDR < 0.05 and s0 > 0.1 to calculate significance. Raw MS data as well as the analysis workflow will be deposited at the ProteomeXchange data repository upon publication.

Immunofluorescence P35 mice were anesthetized by intraperitoneal injection with a lethal dose of 1 μl/g xylazine (VMB Xyl-M 2%), 2 μl/g ketamine (Eurovet Nimatek, 100 mg/ml), and 3 μl/g 0.9% saline. Next, mice were transcardially perfused with 4% PFA, in PBS. Brains were dissected, postfixed in 4% PFA in PBS at 4 °C overnight, washed in PBS and sliced to 50um thick by vibratome. Sections were blocked in 10% NHS, 0.5M Glycine, 0.5% Triton-X 100, 0.02% gelatin in 1X PBS for 2h before incubation with antibodies in 5% NHS, 0.5% Triton-X 100, 0.02% gelatin in 1X PBS with extensive washing with 0.05% Triton-X 100 in 1X PBS between primary and secondary antibodies. Sections were counterstained with DAPI before being mounted on microscope slides with Mowiol-4-88 (Millipore). Imaging was performed on a Slide Scanner Axio Scan.Z1 (Zeiss) or LSM880 with Airyscan (Zeiss).

For spine density analysis, mice were perfused with 4% PFA (EM grade, Science Services), 2% Sucrose in 0.1M PB buffer pH 7.4 and post-fixed 2 h at RT. Brains were sliced at 120um to increase the portion of dendritic tree for each positive neuron.

### Mice injected with turboID AAVs were injected with 2mg/ml biotin in PBS 3h before perfusion

RNAscope in situ hybridization Mice injected with cre-dependent GFP AAV were transcardially perfused at P35 as described. 20um slices were produced on a Vibrating Microtome 7000 (Campden Instruments LTD), mounted on SuperFrost Ultra Plus adhesion slides (Thermo-Fisher) and dried at RT. Sections were stained using the RNAscope™ Multiplex Fluorescent Detection Kit v2 (ACDbio, 323110) following manufacturer protocols. Briefly, sections were baked at 40C for 30 minutes to improve adhesion, treated for 10 minutes with hydrogen peroxide solution and 12 minutes with protease IV washing with 1X PBS in between. RNAscope probes were hybridized at 40C for 2h and subsequently HRP signal developed using OPAL dyes. To counterstain the GFP, sections were blocked for 1h at RT in 5% NHS, 0.3% TritonX-100, in PBS. GFP antibody was diluted in blocking solution 1:500 and added O/N at 4C. after 3 washes in 1X PBS, anti-chicken 488 secondary antibody was added 1:1000 for 1h at RT. After 3 more washes in 1X PBS, sections were counterstained with DAPI for 5 minutes, and coverslipped with Mowiol. Images of GFP positive neurons were acquired on a Zeiss LSM900 using a 20X objective. Same laser power and gain settings were used to image the same gene in the two different mouse lines. Single dots were counted manually.

Dendritic spine imaging and analysis Adult mice were injected in the barrel cortex with 200nl of AAV mix consisting of pAAV-TRE-DIO-FLPo (Addgene #118027), pAAV-TRE-fDIO-GFP-IRES-tTA (Addgene #118026) and either cytoplasmic.turboID, PSD95.turboID or nothing. A week later, mice were injected with 1ml of 2mg/ml biotin in 1X PBS for 3 hours before perfusion with 4% PFA, 2% Sucrose in 0.1M PB buffer pH 7.4 and post-fixed 2 h at RT. Brains were sliced at 120um to increase the portion of dendritic tree for each positive neuron. Brain slices were stained for GFP and biotin. Dendritic spines images were acquired on a Zeiss LSM880 with airyscan detector using a 63X objective and 2.0 zoom. Optimal settings suggested by the software were used for pixel size and optical section size. Laser intensity and gain were adjusted for each image to obtain the best signal to noise ratio for GFP channel. Laser intensity and gain were kept the same for all conditions for the biotin channel. For each brain, 3 dendrites were imaged. Dendritic spines were reconstructed using neurolucida software and classified by morphology.

Human cortex data analysis Processed SNARE-seq2 human cortex dataset was downloaded from Bakken et al.7. The gene expression data consists of 84178 cells selected by Bakken et al. Data analysis was performed using Seurat v4.38. Single-nucleus transcriptomics data was first normalized and highly variable genes were selected using SCTransform (variance stabilizing method) function from Seurat. Principal component analysis was then performed on normalized data. FindNeighbors function was subsequently used to compute the shared nearest neighbor graphs using 35 principal components (chosen using ElbowPlot function in Seurat). Finally, non-linear dimensionality reduction was performed using the uniform manifold approximation and projection (UMAP) method and clusters were obtained using Louvain clustering9 (resolution 0.5). Clusters were then manually annotated based on marker gene expressions.

Layer 5 clusters analysis Cells belonging to layer 5 IT and layer 5 PT clusters were then selected for downstream analysis. Selected cells (5039 cells) gene expression matrix was normalized and scaled using SCTransform function from Seurat v4.38. Differentially expressed genes between cells from layer 5 IT and cells from layer 5 PT clusters were identified by performing Wilcoxon rank sum test using FindMarkers function from Seurat. Genes with adjusted p-value < 0.01 and log fold change > 0.25 were considered as differentially expressed

Software and databases Gene ontology analysis was performed using the R package gprofiler2 (0.2.2) based on gProfiler10. All proteins identified by MS were used as a reference background. Data were plotted using the R package ggplot2 (3.4.2). Data from Sorokina, Mclean, Croning11 were accessed with the R package RSQLite (2.3.1) or directly with MySQLlite software. SYNGO12 database was imported to R from https://syngoportal.org/. Genetrek database was imported to R from https://genetrek.pasteur.fr/. SFARI database was imported to R from https://gene.sfari.org/.

### Antibodies

rat anti HA (Merck, 11867423001, WB: 1:1000, IHC: 1:1000),

Alexa 647-conjugated streptavidin (Thermo Fisher Scientific, S32357, IHC: 1:500),

rabbit Homer1 (Synaptic Systems, 160 003, WB: 1:5000, IHC: 1:1000).

HRP conjugated streptavidin (Thermo Fisher Scientific, SA10001, WB: 1:5000),

mouse anti Gria1 (Millipore, MAB2263, WB: 1:500),

chicken anti-GFP (Aveslabs, GFP-1020, IHC: 1:500),

rabbit anti Tubulin 3 (Abcam, ab18207, WB: 1:2500),

rabbit anti VGLUT2 (Synaptic Systems, 138-403, IHC: 1:5000),

Alexa fluor conjugated secondary antibodies from Invitrogen (1:1000).

## Acknowledgements

We thank Esther Klingler, Patrik Verstreken, Pierre Vanderhaeghen and Anthony Holtmaat for critical reading of the manuscript, and De Wit lab members for helpful discussion and comments. We are grateful to Joris Vandenbempt for mouse colony maintenance and genotyping. We thank Charles Gerfen (National Institute of Mental Health, USA) and Nathaniel Heintz (The Rockefeller University, USA) for generously providing the Tlx3_P56-Cre and Sim1_KJ18-Cre mice. We thank Aya Takeoka (VIB-Imec-KU Leuven NeuroElectronics Research Flanders) for sharing Tlx3-Cre and Rbp4-cre mice. We thank Kristofer Davie (VIB Center for Brain & Disease Research Single-Cell Bioinformatics Expertise Unit) for bioinformatic support. G.M. is supported by Fonds Wetenschappelijk Onderzoek (FWO, Belgium) PhD fellowship 11F1219N and 11F1221N; D.D. is supported by FWO Postdoctoral fellowship 12W5218N, 12W5221N and FWO Research Grant 1513320N; B.L-E. is supported by FWO PhD fellowship 1120821N and 1120823N; L.B. is supported by FWO Postdoctoral fellowship 12ZK221N. J.d.W. is supported by FWO Project Grants G0C4518N, G0A8720N and G0A8320N; FWO EOS Grant G0H2818N; ERANET-NEURON TAO2PATHY G0I3118N and a Methusalem Grant of KU Leuven/Flemish Government.

---

**Extended Data Figure 1.**
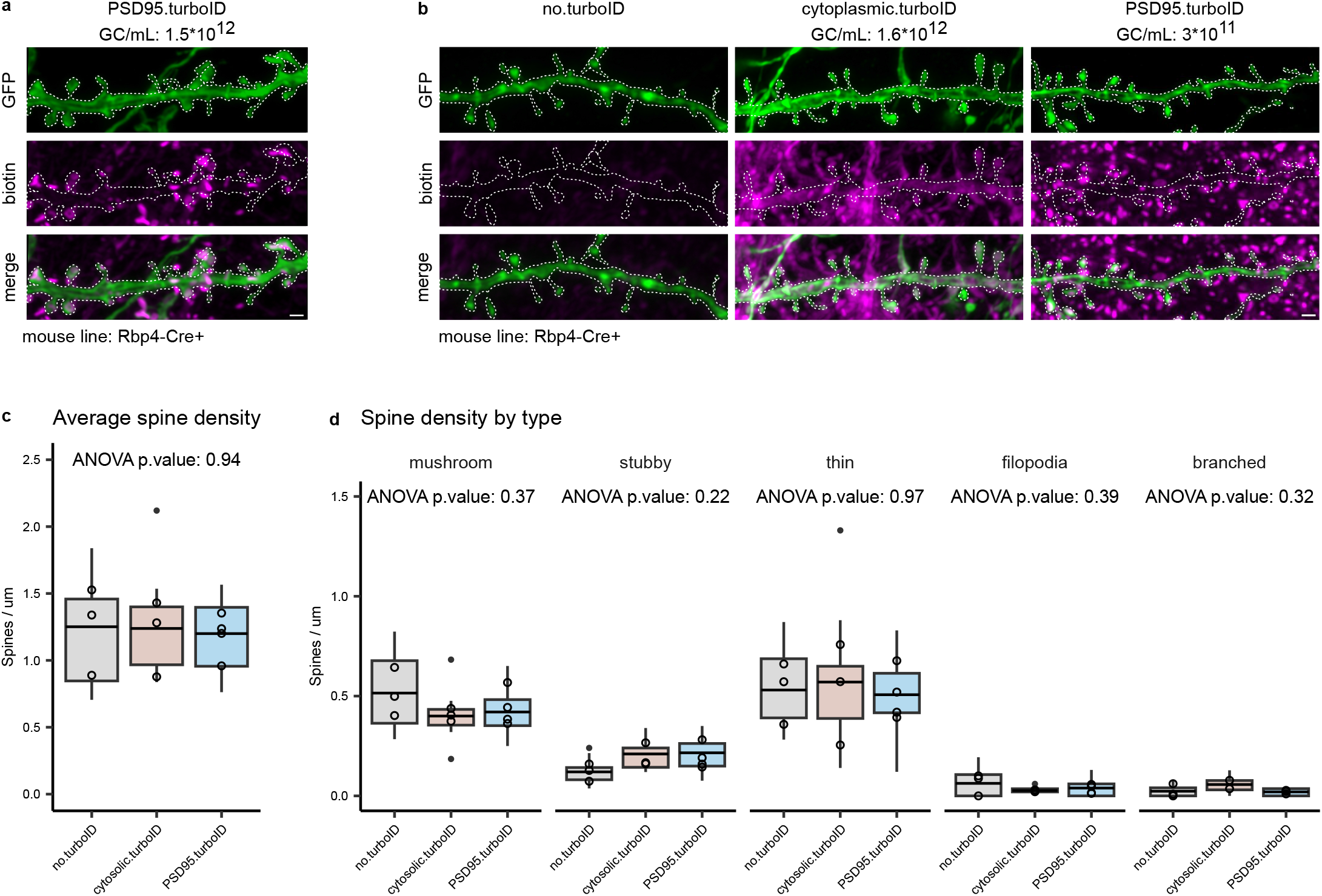
TurboID expression has no effect on dendritic spine density or morphology. **a,** Representative images of dendrites of L5 neurons expressing higher titers of the PSD95.turboID construct affecting spine morphologies. Viral titer is indicated as Genome Copies (GC) per mL. Scalebar is 1 μm. **b,** Representative images of dendrites of L5 neurons expressing different TurboID constructs. Viral titer is indicated as Genome Copies (GC) per mL. Scalebar is 1 μm. **c,** Quantification of spine density. **d,** Quantification of spine density by spine type. N = 3 mice per condition, 2 dendrites per mouse. Boxplots (line across the box describes the mean) show spine density distribution of individual dendrites, empty dots display the average spine density per each replicate. ANOVA statistical analysis is performed on the replicate means.

**Extended Data Figure 2.**
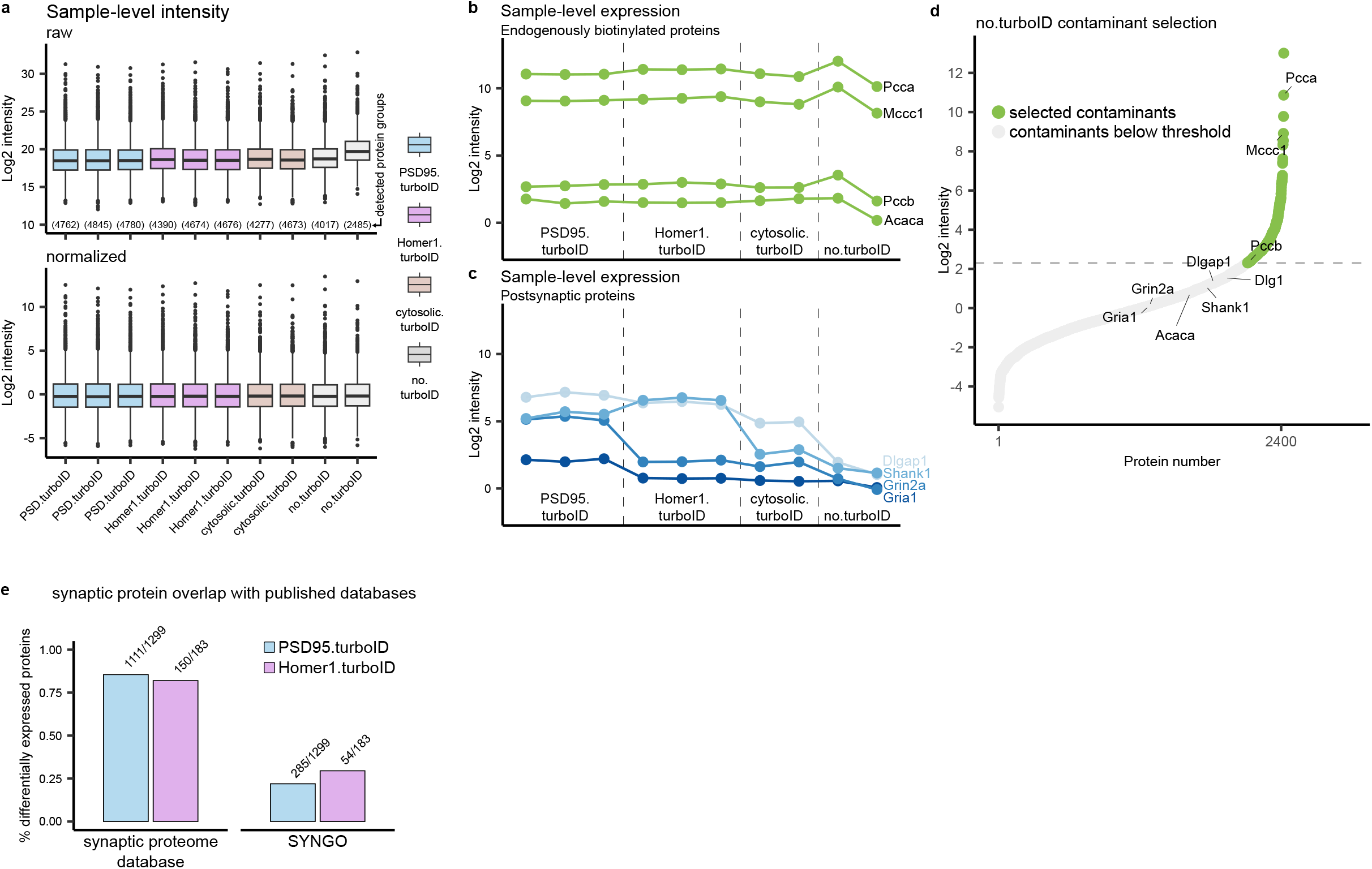
Quality control and analysis for PSD95.turboID and Homer1.turboID synaptic proteome workflow. **a,** Sample-level intensity values for each replicate before and after median normalization. Protein groups detected in each replicate are reported on the top graph. **b,** Intensity levels in each sample for 4 endogenously biotinylated proteins. **c,** Intensity levels in each sample for 4 known PSD proteins. **d,** Proteins detected in all 4 no.turboID replicates are ranked by intensity. Endogenously biotinylated proteins and known PSD proteins are labelled. Threshold for selecting contaminants is based on the expression of endogenously biotinylated proteins and known PSD components. **e,** Overlap of the enriched proteins against the synaptic proteome database and SYNGO. See Supplementary Table 1 for complete annotation.

**Extended Data Figure 3.**
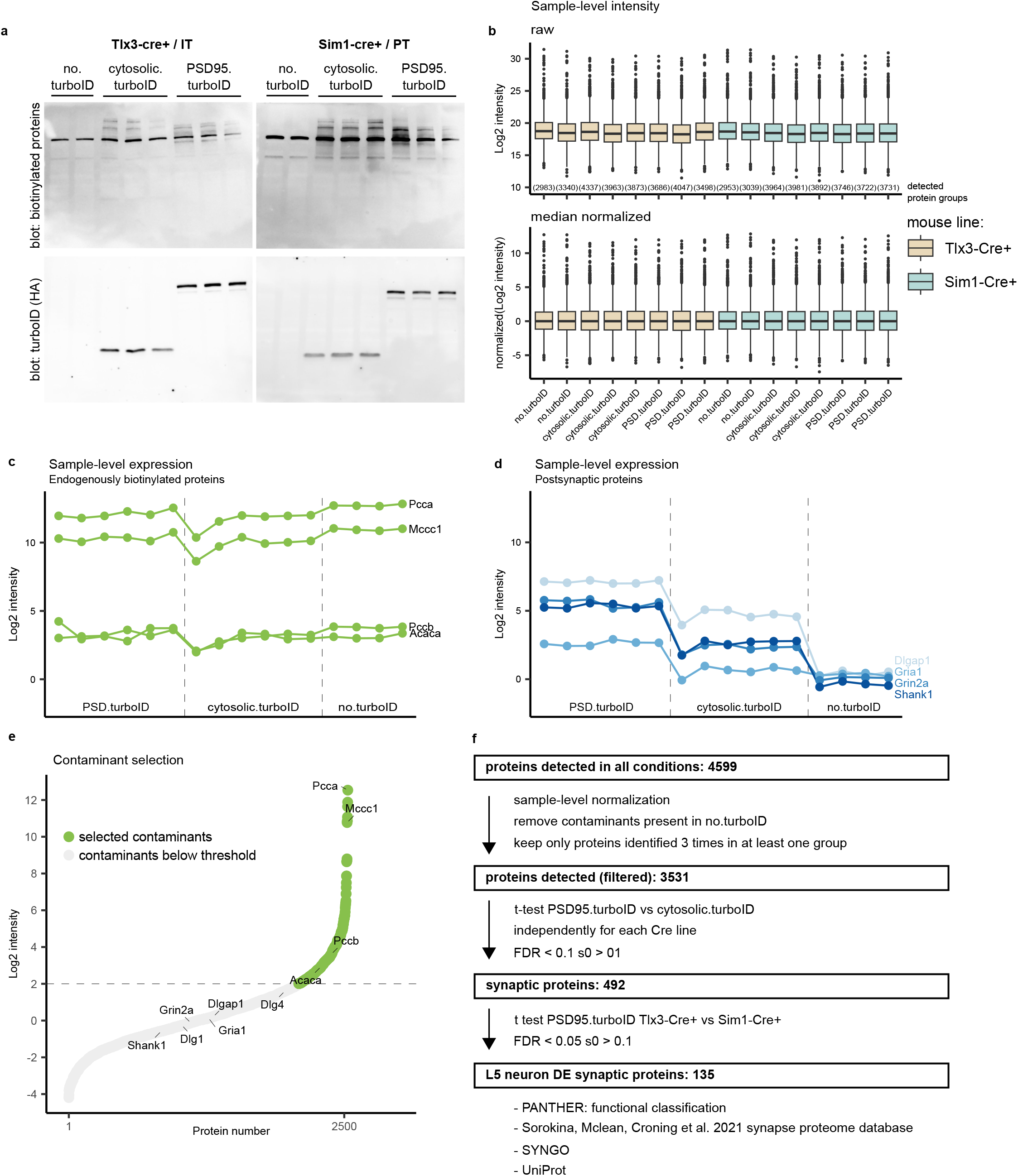
Quality control and analysis for cell type-specific TurboID synaptic proteome workflow. **a,** Western blots for biotinylated proteins and TurboID-HA on replicates confirm successful biotinylation in every sample and correct TurboID construct expression. **b,** Sample-level intensity values for each replicate before and after median normalization. Protein groups detected in each replicate are reported on the top graph. **c,** Intensity levels in each sample for 4 endogenously biotinylated proteins. **d,** Intensity levels in each sample for 4 known PSD proteins. **e,** Proteins detected in all 4 no.turboID replicates are ranked by intensity. Endogenously biotinylated proteins and known PSD proteins are labelled. Threshold for selecting contaminants is based on the expression of endogenously biotinylated proteins and known PSD components. **f,** Proteomic analysis workflow to enrich for synaptic proteins and select candidates from differentially expressed proteins.

**Extended Data Figure 4.**
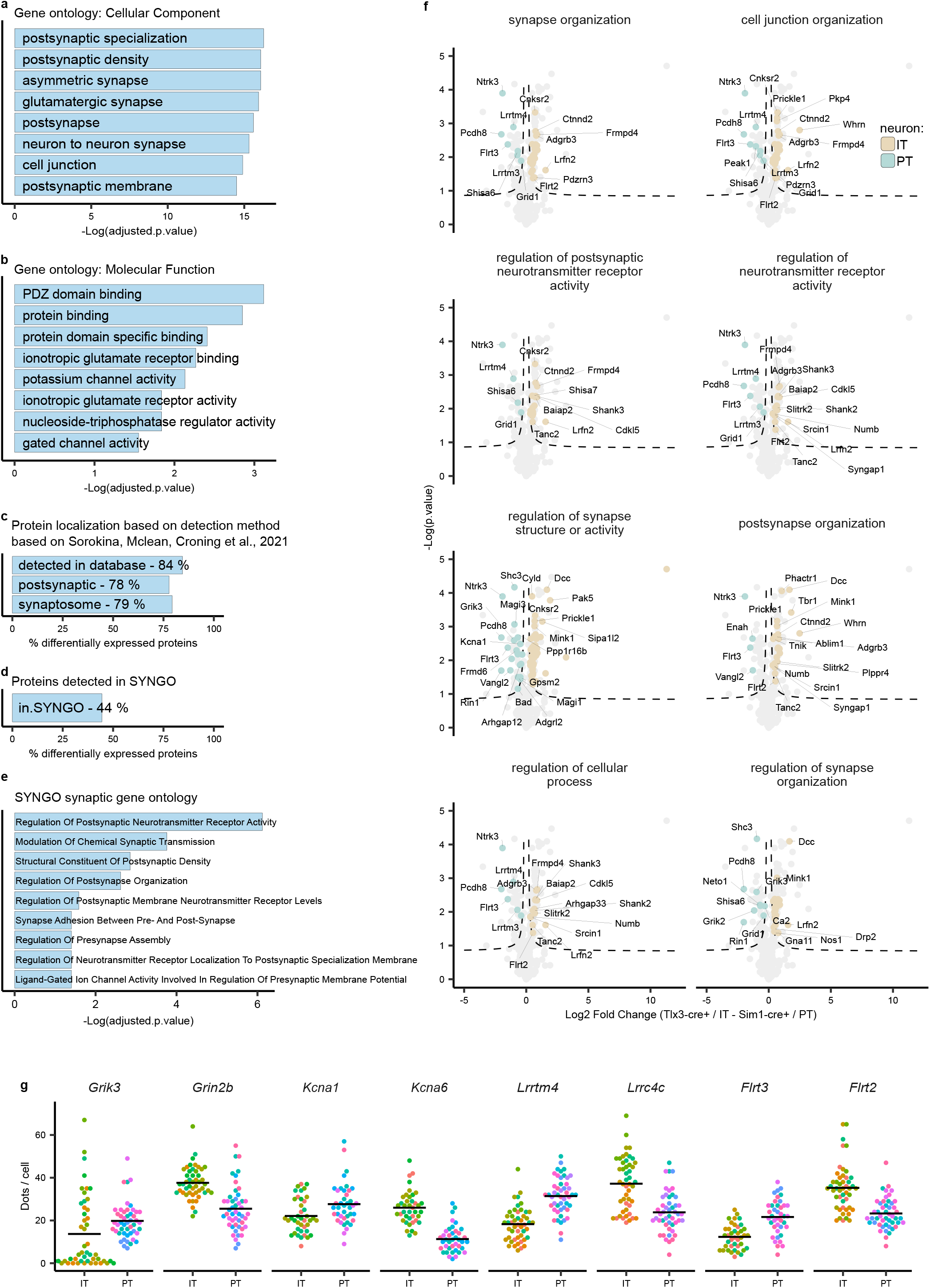
Gene ontology analysis of differentially expressed proteins in layer 5 neurons. **a,** Gene ontology for Cellular Compartment and **b,** Molecular Function categories using differentially expressed proteins as input. **c,** Analysis of differentially expressed proteins using the synapse proteome database^63^. 85% were detected in the database. 79% and 82% were identified in postsynaptic fractionation studies and synaptosome fractionation studies, respectively. **d,** 46% of differentially expressed proteins were detected in the SYNGO database^46^. **e,** SYNGO gene ontologies. **f,** Annotation of proteins found in the Biological Process gene ontology terms (Figure 3b). Not all protein names are displayed due to space limitations, see Supplementary Table 2 for complete annotation. **g,** Plots from Figure 4f and 4h showing distribution of the analyzed cells. Each color represents a different mouse, each dot represents a single cell.

